# Acute toxicity of Leaf extracts of *Enydra fluctuans* Lour in Zebrafish (*Danio rerio* Hamilton)

**DOI:** 10.1101/2020.03.27.012021

**Authors:** Jobi Xavier, Kshetrimayum Kripasana

## Abstract

The present study was focused on the concentration dependent changes in oral acute toxicity of leaf extracts of *E. fluctuans* in zebrafish. The study was also aimed at the details of histopathological changes in gill, liver, brain and intestine of zebrafish exposed to the leaf extracts of the plant *E. fluctuans*. *Enydra fluctuans* Lour is an edible semi-aquatic herbaceous plant used widely for the alleviation of the different diseases. Since there were no toxicity studies conducted on this plant the present study was an attempt to look into the elements of toxicity of the plants. Two types of experiments are conducted in the present study. First, the acute oral toxicity study was conducted as per the OECD guidelines 203. Second, histopathological changes were observed in the fishes exposed to the lethal concentrations of plant extract. The oral acute toxicity studies conducted on Zebrafish have revealed that the leave extracts of *E. fluctuans* were toxic to the tested fish at the concentration of 200 mg/kg body weight. The histopathological studies conducted on intestine of treated fishes showed that treatment has induced rupturing of villi structure and fusion of villi membrane and detachment of villi structure from basal membrane of intestine. The histology of the liver also showed severe vacuolization in the cells while it is not affected in control. The studies on gills showed the detachment of basal epithelial membrane in gills compared to control which might have led to death of the fish. The histopathological observations of brain tissues treated with test samples also revealed the marked impingement in brain parenchyma while the control is normal without impingement of brain.

## 1. Introduction

With the development of herbal industry, people started to use many plant extracts in the preparations of herbal products commercially. People are of the assumption that all plant products are safe to consume and there is no need to investigate on their safety aspects. Many studies have not been conducted on the toxicity of those herbal products. The present study was designed to investigate on toxicity level of leaf extracts of the plant *Enydra fluctuans* Lour in the zebrafish. The toxic agent is mostly released from sources like leaf, fruits and barks of plants, animals, and microorganisms. As a toxic agent, it will transmit the toxic substance through the various modes of transmission mainly via direct contact. A toxicology test is necessary, not only for allopathic medicine but also for complementary and alternative medicine to discover any adverse effects which are not known until the signs and symptoms develop upon high consumption [1]. The Assessment of toxicity using acute toxicity bioassay can prove the safety of traditional medicine using the plants *E. fluctuans* and promote its consumption.

The importance of toxicity testing is to provide dose dependent changes against the toxicity effect, study the safety of components in the sample, and authenticate methods of investigating toxicity [2]. In the toxicity studies, Zebrafsh (Danio rerio) is used as a good model to study the toxicity effect. It is used to inspect the bioactive compounds of the sample through toxicity assay. Commonly used mammalian models have some drawbacks with higher cost and prolonged time for results, being yet ethically questionable [1, 3, 4]. Studies have proved that humans show genetically great similarities of genomic sequences and brain patterning with the Danio rerio. Thus, this makes zebrafish an advantageous assay in exploring many diversions of toxicology study yielding a prompt outcome [4–6]

### Enydra fluctuans

Lour [Figure 1] is an edible semi-aquatic herbaceous plant belonging to the family Asteraceae. The vernacular name of this plant is Helencha, Hinchashak, Harhach and the English name Water Cress, Marsh Herb etc [7]. The leaves are slightly bitter and the leaves are used to cure inflammation, skin diseases, laxative, bronchitis, nervous disorders, leucoderma, biliousness and small pox. It possesses biological value and its fuel extract has been reported to own analgesic, cytotoxic, antimicrobial, hepatoprotective, hypotensive, CNS depressant, and antidiarrheal activity [8].

**Figure 1:**
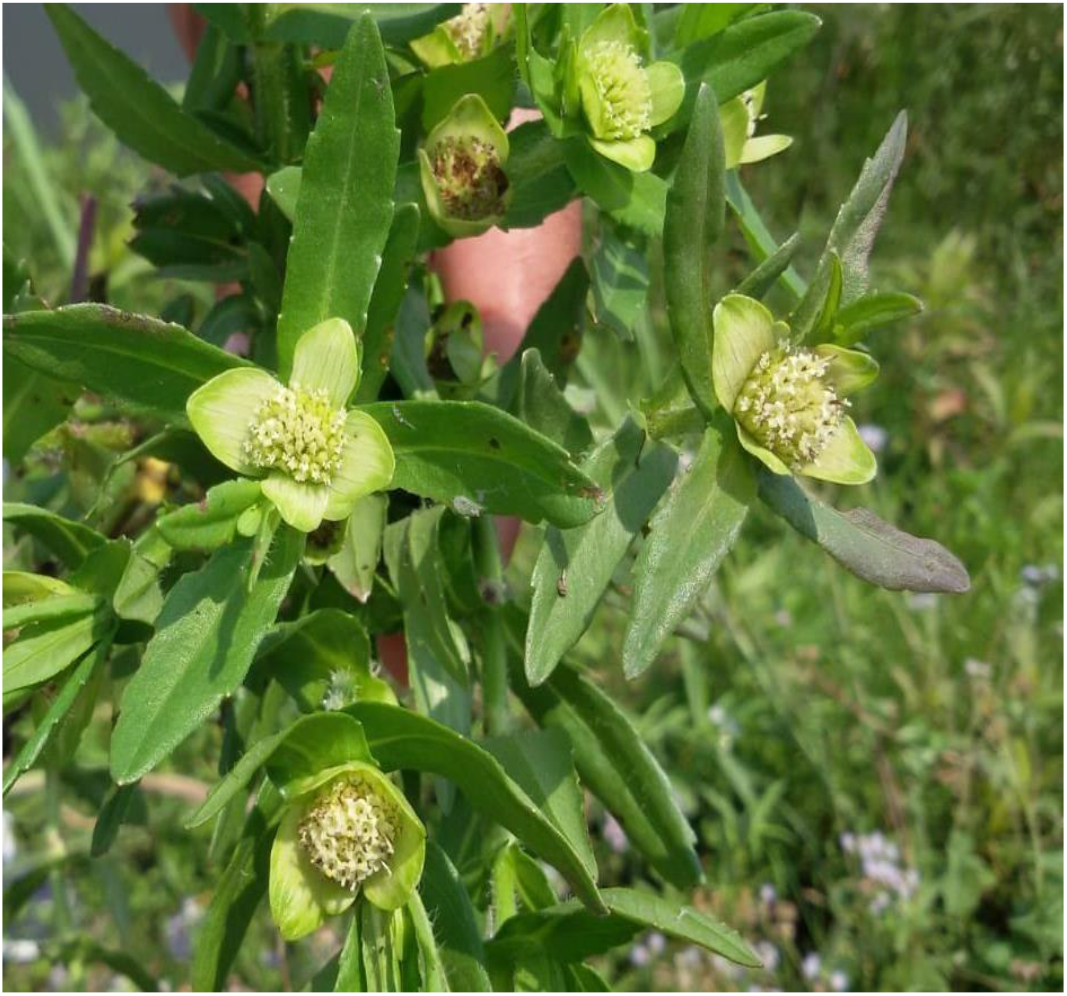
Enydra fluctuans

The objectives of the present study were to investigate on concentration dependent changes in oral acute toxicity of leaf extracts of *E. fluctuans* in zebrafish. The study was also focused on the details of histopathological changes in gill, liver, brain and intestine of zebrafish exposed to the leaf extracts of the plant *E. fluctuans*. Histopathological evaluation is one of the important parts of the assessment of the adverse effects of the plant extracts on the whole organism. Plants are not always safe to consume as they may contain poisonous chemical compounds in the different parts of the plant. Many studies have revealed that the many plant extracts become toxic to the different organs on concentration dependent manner and can lead to lethality. Many of the herbal products that are used as remedy to different diseases are not undertaken for their toxicity level [8]. The present study was an attempt to understand the toxic level of widely used plant *E. fluctuans*. In the present study, the hsitopathological studies were performed in fishes which showed the mortality. It is hypothesized that higher concentrations of the leaf extracts could be detrimental to the fish exposed to the leaf extracts and affect the susceptible organs which can be examined histopathologically. Since there are no published data on the histopathological evaluations of the fish exposed to the different concentrations of the leaf extracts of *E. fluctuans*, the present study has been undertaken.

## 2. Materials and Methods

### 2.1 Preparation of ethanolic leaf extract of *Enydra fluctuans*

*Enydra fluctuans* (Authentication/PDDUIAS/2019/01) was collected from Yaiskul of Imphal West of Manipur, India. The leaves of the plants were dried and ground into fine powder. The extracts of the plants were prepared using 5 g of dried powder in 50 mL of the ethanol. The extracts were filtered using rotary evaporator and various concentrations of the test solutions were prepared.

### 2.2 Oral Acute toxicity of ethanolic leaf extract of *Enydra fluctuans*

Acute toxicity of ethanolic leaf extract of plant samples *Enydra fluctuans* was tested for acute toxicity in zebra fish model (*Danio rerio*-IAEC/CU/ZF/2018/01) as per the OECD guidelines 203 [9]. The fish were exposed to the test substance preferably for a period of 96 hours. Mortalities are recorded at 24, 48 and 96 hours and the concentrations which kill 50 percent of the fish were recorded.

The fish were inspected after 24, 48, 72 and 96 hours. Concentrations of 12.5, 25, 50, 100, 200, and 400mg/L were selected as effective concentrations for performing the main toxicity tests of the plant extracts of different concentrations. The fish were exposed to the sample based on a static exposure regime. For every experiment, 7 healthy fishes were directly transferred into each prepared concentration. Control groups (7 fishes) were also included for each treatment. The mortalities were recorded at 24, 48, 72 and 96 hours post exposure and the LD50 values were calculated [8]. Fish were considered dead if there is no visible movement and upon touching of the caudal peduncle produces no reaction. Dead fishes were removed when mortalities are recorded. LD 50 was determined based on the concentration of the test substance in water which killed 50 percent of a test batch of fish within a particular period of exposure was observed.

### 2.3 Statistical Analysis

The median lethal concentration (LC_50_) of the acute toxicity experiment was calculated from the data using the PROBIT function as described by Finney in 1971 [10] and analyzed by IBM SPSS Statistics 21.0 software with 95% confidence limits. The safe level estimation after 96 hr exposure of zebra fish to the leaf extracts of *E. fluctuans* was carried out based on Hart et al. [11], Sprague [12], Committee on Water Quality Criteria (CWQC, 1972) [13], National Academy of Sciences/National Academy of Engineering (NAS/NAE, 1973) [14], and International Joint Commission (IJC, 1977) [15].

### 2.4 Histopathology of Zebrafish based on Acute Toxicity

Fish were euthanized after cold temperature anesthesia below 15° C and Liver, Heart, Brain and intestine tissues were removed and smear stained with Haematoxylin and Eosin using standard protocols. For behavioral and motor toxicity swim motion test and dive tank test were performed. The gills and skin of experimental fish were collected after 96hr exposure period and fixed in 10% neutral buffered formalin, and paraffin, sectioned at 5μm and stained with Haematoxylin and Eosin.

## 3. Results

### 3.1 Oral Toxicity studies of ethanolic extracts of the Leaf of *Enydra fluctuans* in *Danio Rerio*

The study conducted on acute oral toxicity revealed that the leaf extracts of *Enydra fluctuans* were more toxic to *Danio rerio*. Medial lethal concentration (LC_50_) is considered the most accepted basis to determine the acute toxicity test. In this toxicity study, 100 % mortality was observed which clearly indicated the toxicity of the plant extract. The LC_50_ values with 95% confidence intervals of different concentrations of the leaf extracts of *E. fluctuans* were 204.132, 170.513, 139.478 and 92.956 mg/L for 24, 48, 72 and 96 hrs, respectively [Table 1]. Concentration dependent change in the mortality during 24, 48, 72 and 96 hrs exposure to leaf extract for the plant sample was observed in the fishes experimented. The plant extract was found to be lethal on the first day of the experiment within 24hr at the highest concentrations of 400mg/L since it killed all the seven fishes in the experiment. A dose dependent increase and the time dependent decrease were observed in mortality rate as the exposure time increased from 24 to 96 hrs, the median lethal concentration was reduced. A negative perfect correlation was observed between the exposure time and the LC_50_ [Table 2].

**Table 1.**
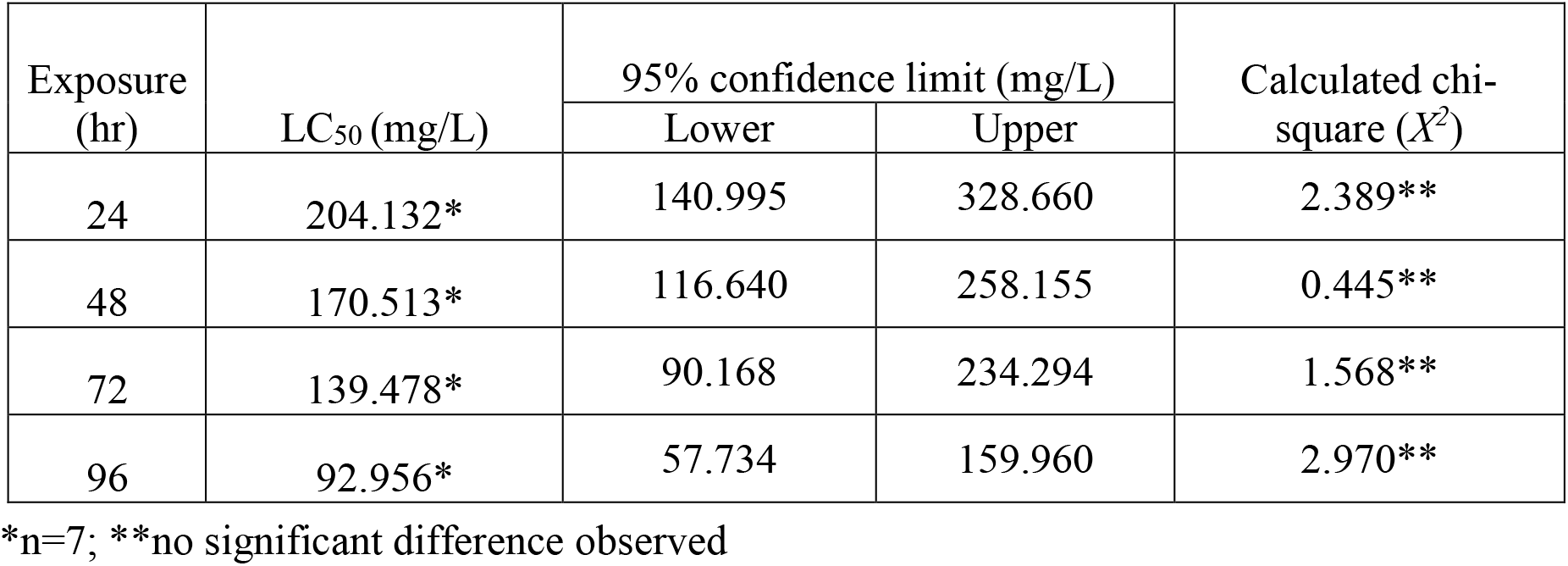
Lethal Concentration (LC_50_) of Leaf extracts of *Enydra fluctuans* with 95% confidence limit depending on exposure time (hr)

| Exposure (hr) | LC<sub>50</sub> (mg/L) | 95% confidence limit (mg/L) |  | Calculated chi-square (X<sup>2</sup>) |
| --- | --- | --- | --- | --- |
|  |  | Lower | Upper |  |
| 24 | 204.132* | 140.995 | 328.660 | 2.389** |
| 48 | 170.513* | 116.640 | 258.155 | 0.445** |
| 72 | 139.478* | 90.168 | 234.294 | 1.568** |
| 96 | 92.956* | 57.734 | 159.960 | 2.970** |

\*n=7; \*\*no significant difference observed

**Table 2.** Correlation Analysis among between exposure time and LC_50_

|  |  | LC<sub>50</sub> | Exposure time |
| --- | --- | --- | --- |
| LC<sub>50</sub> | Pearson Correlation | 1 | -.966* |
|  | Sig. (2-tailed) |  | .034 |
|  | N | 4 | 4 |
| Exposure Time | Pearson Correlation | -.966* | 1 |
|  | Sig. (2-tailed) | .034 |  |
|  | N | 4 | 4 |

\*. Correlation is significant at the 0.05 level (2-tailed).

Fish were considered dead if there is no visible movement and if touching of the caudal peduncle produces no reaction. Dead fishes were removed when mortalities are recorded. LD50 was determined based on the concentration of the test substance in water which killed 50 percent of a test batch of fish within a particular period of exposure. Toxic effects of this plant sample of *Enydra fluctuans* were observed. At dose of 200mg/L >70% mortality was observed in zebra fish. Mortality was not observed at 50mg/L of plant extract. Lethal dose to kill 50% of test fishes were recorded as 400mg/L, 200mg/L at 24hr and 96hr exposure period respectively. The results indicated that the ethanolic extract did not induce toxicity when used the dose of ≤ 50mg/L [Table 1]. No mortality was observed in the control during the experimental period. Variations were observed in safe levels estimated by different methods at 96 hrs of exposure of zebra fish to the leaf extracts of the plant as shown in Table 3.

**Table 3.**
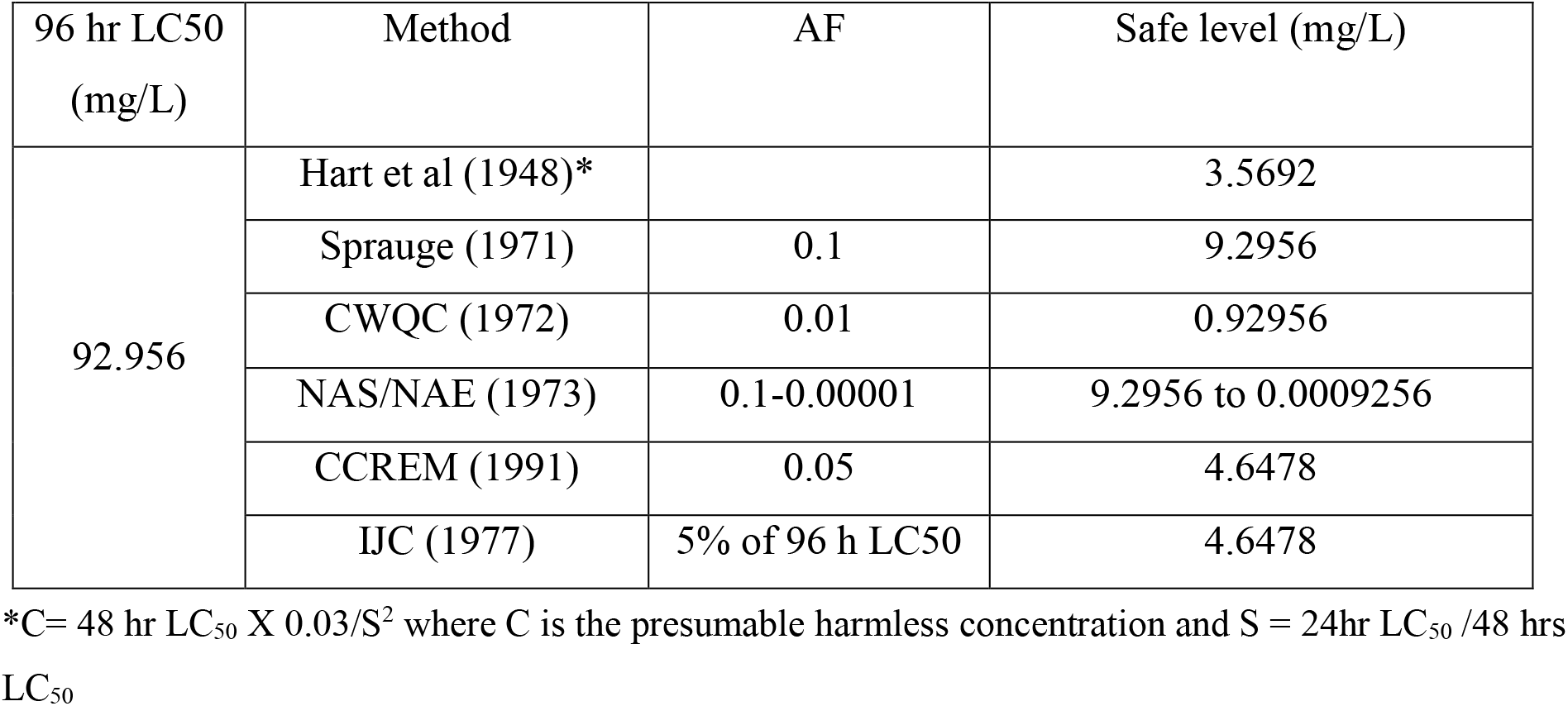
Estimation of safe levels of the plant extracts at 96 hr of exposure

### 3.2 Histopathology of Treated Zebrafish of *Enydra fluctuans* plant extract with Control

Treatment of the zebrafish with plant extracts induced the rupturing of villi structure and fusion of villi membrane and detachment of villi structure from basal membrane in the intestine [Figure 2 & 3] Tissues treated with samples have shown detachment of basal epithelial membrane in gills compared to control [Figure 4 & 5]. Liver tissue in treated samples also showed severe vacuolization in the cells [Figure 6 & 7]. Brain tissue treated with test samples was marked with impingement in brain Parenchyma [Figure 8 & 9].

**Figure 2.**
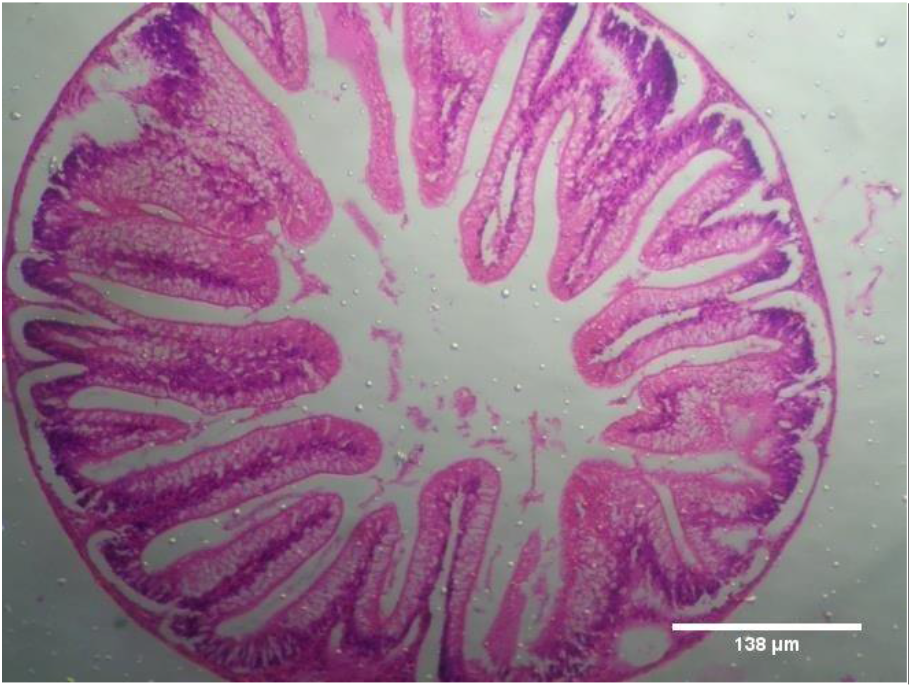
Control- Intestine

**Figure 3.**
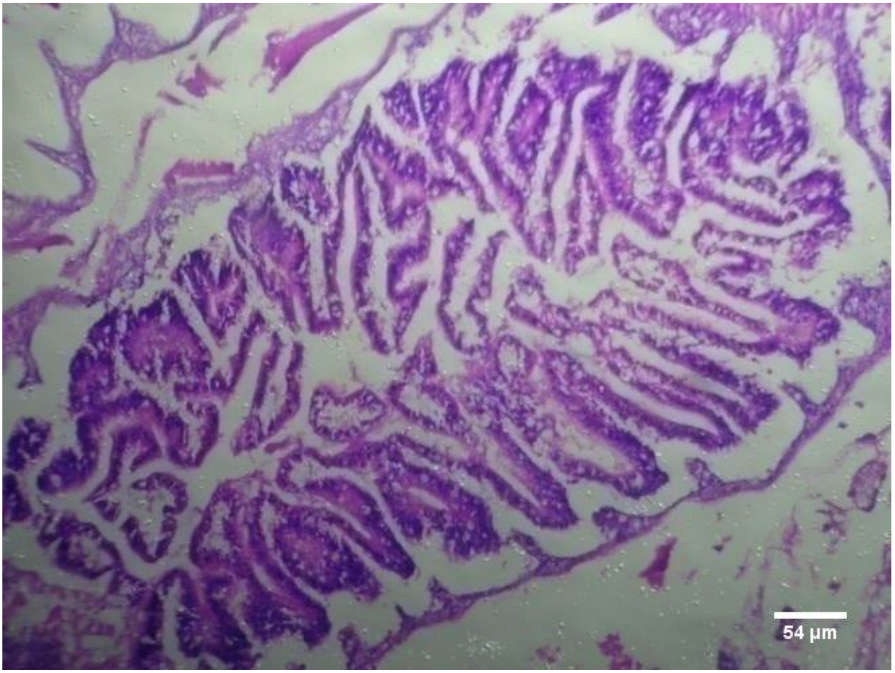
Intestine treated with leaf extract

**Figure 4.**
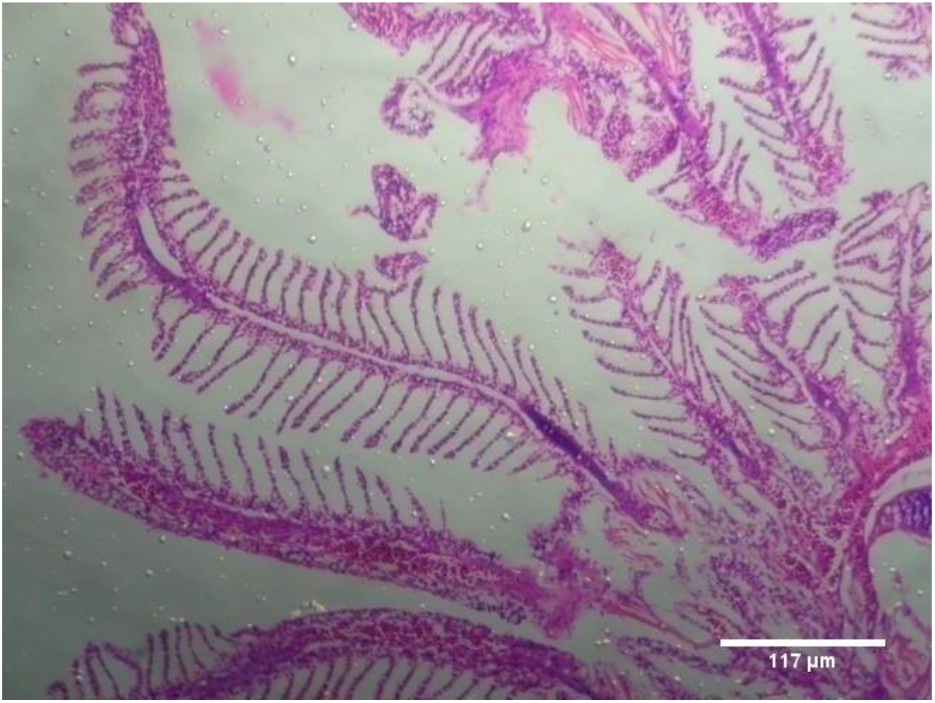
CONTROL- GILL

**Figure 5.**
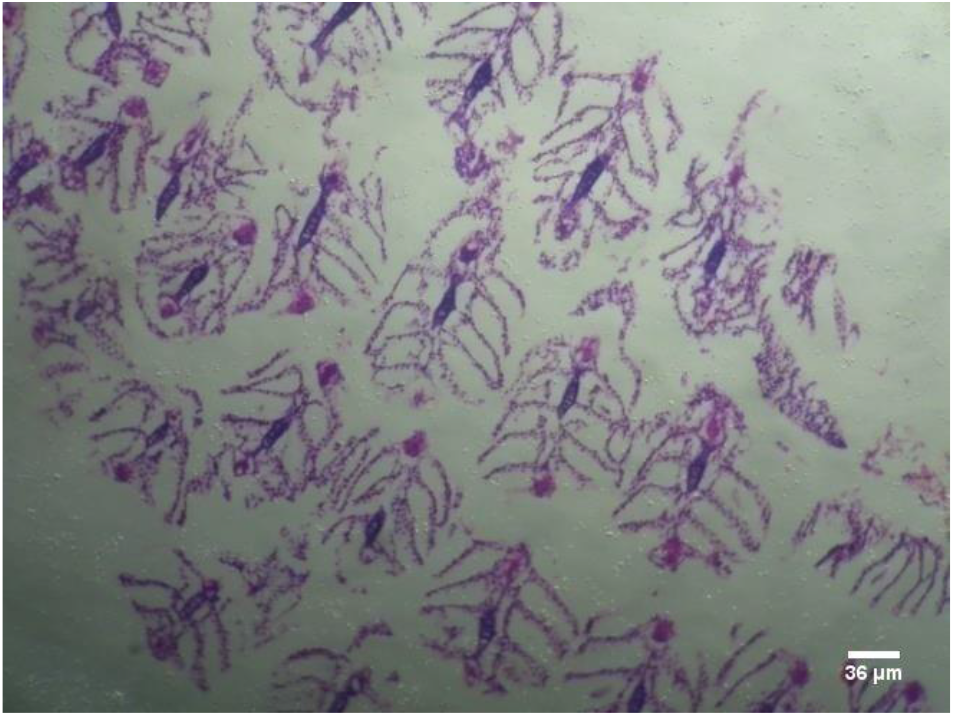
Gill treated with leaf extract

**Figure 6.**
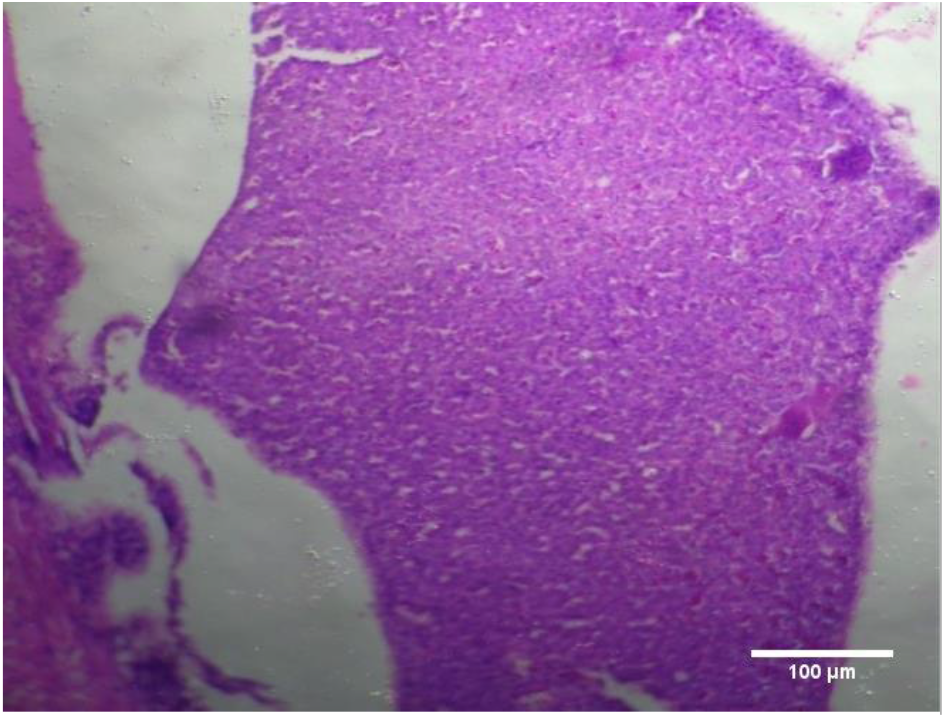
CONTROL- LIVER

**Figure 7.**
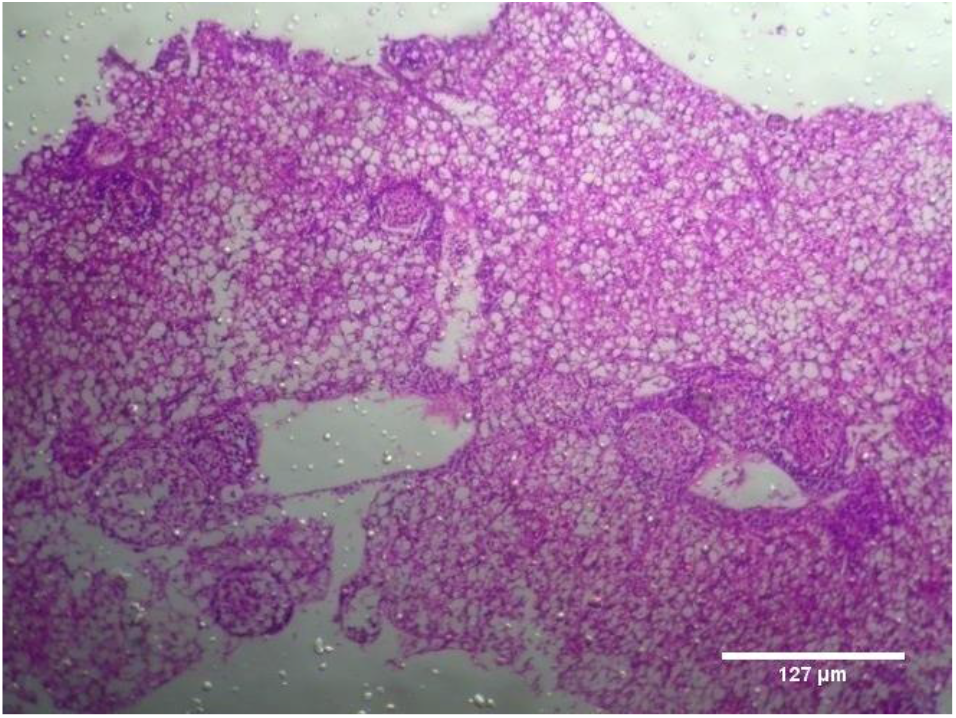
Liver treated with leaf extract

**Figure 8.**
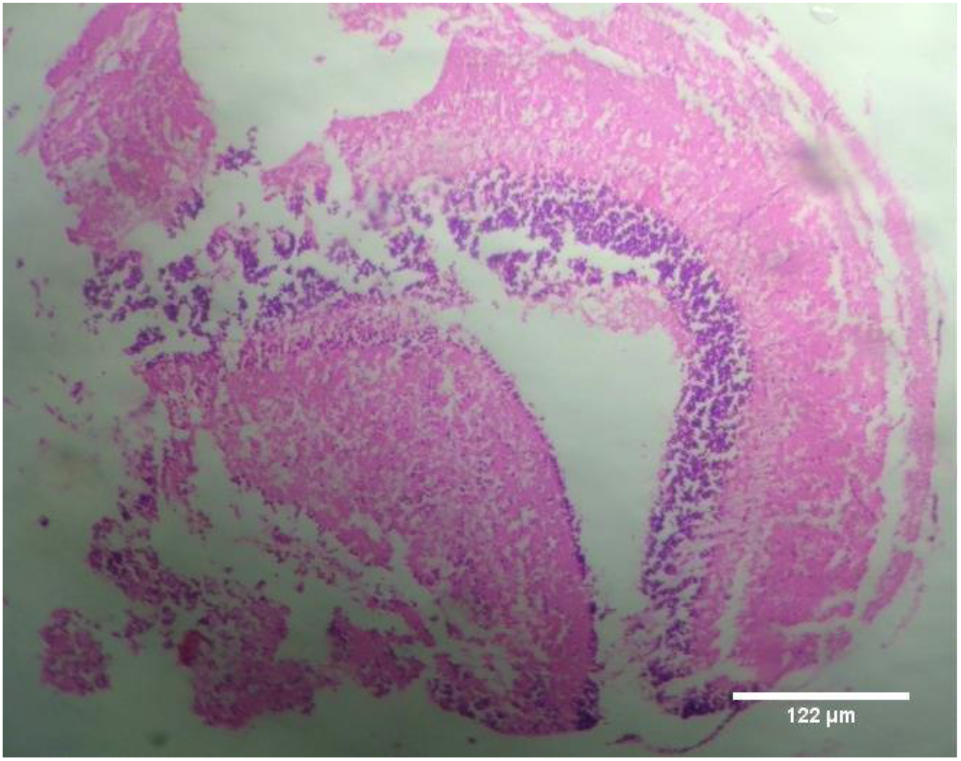
CONTROL- BRAIN

**Figure 9. -.**
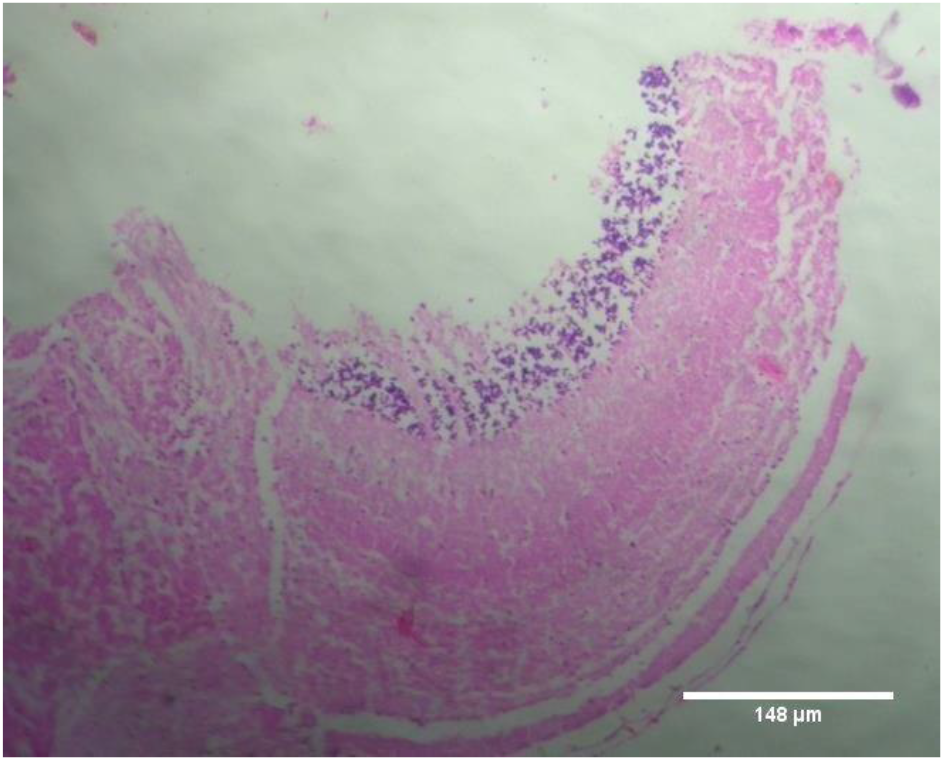
Brain treated with leaf extract

**Fig 2- 9. Histopathology of Intestine, Liver, Gills and Brain from Zebrafish expose to 400mg/L**

### 3.3 Behavioural responses

The test fish *Danio rerio* exposed to various lethal concentrations of leaf extracts of *E. fluctuans* exhibited altered behavioral responses. It was observed that in lethal concentrations of plant extracts, hyperactive behavior was increased initially and subsequently reduced expressing the sign of distress. The body showed depigmentation along with mucus secretion. Reduced swimming performance lethargic movements were observed in the fish exposed to lethal concentrations of the plant extracts.

## 3. Discussion

In this study, we analyzed the sensitivity of the leaf extracts of *E. fluctuans* in the experimental fish and have shown that the lethal effect of the leaf extract was highest on zebrafish at the exposure time of 96 hrs. The LC_50_ at 96 hrs of exposure of zebra fish in the present study was 2.798, which was lower than the LC_50_ of tests conducted with cd^2+^ and Zn^2+^ in zebrafish i.e., 6.5 and 44.48 mg/L and greater than the 96 hr LC_50_ with Cu^2+^ and Hg^2+^, i.e., 0.17 and 0.14 mg/L [16]. Studies conducted by Nwani *et al* [17] on the fish *Channa punctatus* have revealed the toxicity and expressed in terms of LC_50_ for glyphosate and atrazine as 32.540 and 42.380 mg/L and the 96 hr LC_50_ with carbosulfan, as 0.268 mg/L. Acute oral toxicity studies of *Momordica charantia* in zebrafish also reported the lethality of the fish at the concentrations of 50mg/kg bw of the plant extracts [8].

Histopathology of zebrafish treated with the plant extracts and control was studied in the present study. Histopathological studies conducted on gills, intestines, liver and brain isolated from the treated fish showed abnormal conditions when compared with the control. The histopathological studies conducted on intestine of treated fishes showed that treatment has induced rupturing of villi structure and fusion of villi membrane and detachment of villi structure from basal membrane of intestine, which lead to the death of the fish. Intestinal villi (singular: villus) are small, finger-like projections that extend into the lumen of the intestine, Villi increases the internal surface area of the intestinal walls making available a greater surface area for absorption. The studies on gills showed the detachment of basal epithelial membrane in gills compared to control which might have lead to death of the fish. Several studies have been made in the past on the histological organization and physiology of gills in a variety of fishes [18]. Gills are very important for the fishes since they are the main site of gaseous exchange and involved in osmoregulation [19]. The studies conducted by Rajini et al (2015) have revealed that the structural damages of gill like inflammatory cell infiltration, minimal congestion in primary lamellae, fusion of secondary lamellae, diffused epithelial hyperplasia and multifoci mucus cell hyperplasia were observed in the zebrafish exposed to the sub-lethal concentrations of combination pesticides [20].

The present study conducted on liver also showed severe vacuolization in the cells while it is not affected in control. Several studies on histopathology of liver revealed that the swelling with non-lipid cytoplasmic vacuolation of diffusely distributed hepatocytes were seen consistently after mild acute and subacute liver injury. Rajini et al reported that moderate, diffused to severe cytoplasmic vacuolation, diffused minimal to mild sinusoidal congestion, steatosis, pyknotic, karyorrhectic nuclei with complete dissolution of necrotic hepatocytes were observed in the liver of fishes exposed to pesticides [21]. Severe necrotic liver damage was observed in the fish treated with the extracts of *M. charantia* and was considered the cause of damage, which happens due to the lethal doses of momordin [8]. Momordin is found to be one of the major phytocompounds present in the *Momordica charantia, which* has pivotal role in the induction of toxicity in animals and fishes [8, 22]. Several lines of evidence point to the possibility that this change may reflect a cellular adaptation beneficial to the host, rather than a degenerative change. But in the present study since fishes exposed to toxic level were found to be dead those cellular degenerations were clear evidence of toxicity. The histopathological observations of brain tissues treated with test samples also revealed the marked impingement in brain parenchyma while the control is normal without impingement of brain. Histological investigation of the brain of *Channa punctatus* in response to acute and subchronic exposure to the pesticide Chlorpyrifos revealed detachment in the superficial zone of the Stratum opticum, Stratum marginal due to degeneration of neuronal cells, spongiosis, congestion, necrosis and appearance of clear areas around the nucleus of mononuclear cells in the lining of the Stratum fibrosum griseum superficiale, Stratum griseum centrale, and Stratum album central [23].

## 4. Conclusion

In view of the data of the present study, it can be concluded that the leaf extracts of *Enydra fluctuans* is toxic to the fishes in a concentration dependent manner and thus can be toxic to human beings at the higher concentrations. So that dosage dependent use of the plant is very much required to avoid toxicity. Lower concentrations of the plant extract are found to be safe and can be consumed in lower concentrations. It is also reported that the histopathological changes in fishes exposed to 200mg/L are more severe and to be very cautious of the use of the plant in daily life.

## Data Availability

The data used to support the findings of this study are available from the corresponding author upon request.

## Conflicts of Interest

The authors declare that there are no conflicts of interest.

## Authors’ Contributions

All the listed authors have made a significant scientific contribution to the research in the manuscript. Both authors performed the experiment, analysed the data, and wrote the manuscript.

## Acknowledgments

This current study was supported by CHRIST (Deemed to be University) and the authors are grateful to the management and staff of CHRIST (Deemed to be University), Bengaluru.

